# Host cell-specific metabolism of linoleic acid controls *Toxoplasma gondii* growth in cell culture

**DOI:** 10.1101/2024.03.22.586332

**Authors:** Nicole D. Hryckowian, Caitlin Zinda, Sung Chul Park, Martin T. Kelty, Laura J. Knoll

## Abstract

The obligate intracellular parasite *Toxoplasma gondii* can infect and replicate in any warm-blooded cell tested to date, but much of our knowledge about *T. gondii* cell biology comes from just one host cell type: human foreskin fibroblasts (HFFs). To expand our knowledge of host-parasite lipid interactions, we studied *T. gondii* in intestinal epithelial cells, the first site of host-parasite contact following oral infection and the exclusive site of parasite sexual development in feline hosts. We found that highly metabolic Caco-2 cells are permissive to *T. gondii* growth even when treated with high levels of linoleic acid (LA), a polyunsaturated fatty acid (PUFA) that kills parasites in HFFs. Caco-2 cells appear to sequester LA away from the parasite, preventing membrane disruptions and lipotoxicity that characterize LA-induced parasite death in HFFs. Our work is an important step toward understanding host-parasite interactions in feline intestinal epithelial cells, an understudied but important cell type in the *T. gondii* life cycle.

## Introduction

*Toxoplasma gondii* is the causative agent of toxoplasmosis, a potentially devastating disease for immunocompromised people (1, 2). As an obligate intracellular parasite, *T. gondii* must adapt to a wide variety of cell types as it moves through the tissues of its warm-blooded hosts. Intestinal epithelial cells are the first host cell type that *T. gondii* encounters following oral infection. Little is known about the biology of *T. gondii* infection in the intestine, despite it being the gateway to dissemination throughout the body and serving as the site of parasite sexual development in feline hosts (3, 4, 5).

The intestinal epithelium comprises many cell types, including absorptive cells, secretory cells, stem cells, and patrolling immune cells. Approximately 90% are enterocytes, which are absorptive cells whose primary function is to uptake, package, and transport nutrients like amino acids, sugars, and lipids from the intestinal lumen into circulation (6, 7, 8). While the enterocyte-dominated intestinal epithelium is a transient phase of *T. gondii* infection in intermediate hosts (9, 10), enterocytes are an essential part of the parasite’s sexual life cycle because feline small intestinal enterocytes are the sole site of sexual development (3). Thus, understanding how *T. gondii* survives and replicates inside nutrient-rich gut cells is important for understanding how it completes its life cycle.

Enterocytes efficiently absorb and transport numerous lipids into the blood, including fatty acids, phospholipids, cholesterol, and acylglycerols (7). Polyunsaturated fatty acids (PUFAs) are an important subclass of fatty acids that serve as structural components of membranes, sources of energy via fatty acid beta-oxidation, precursors to bioactive lipids like eicosanoids, and as ligands for metabolic and immune receptors like peroxisome proliferator-activated receptor (PPARs) (11, 12, 13, 14). Some PUFAs also show *vitro* histone deacetylase (HDAC) inhibitory activity, suggesting they may epigenetically influence gene expression by influencing chromatin accessibility (15). Mammals are unable to synthesize two 18-carbon PUFAs that they must obtain from dietary sources: the *n*-3 PUFA α-linolenic acid (ALA, 18:3) and the *n*-6 PUFA linoleic acid (LA, 18:2) (16, 17).

Felines have further dietary requirements for the desaturation and elongation products of ALA and LA because they lack intestinal delta-6-desaturase activity (18, 19). Our lab previously showed that mimicking cat-like LA accumulation in intestinal organoid-derived monolayers causes *T. gondii* to begin sexual development (20). However, we still do not understand mechanistically how LA induces *T. gondii* sexual development or how other species- and cell type-specific aspects of cat intestinal lipid metabolism influence *T. gondii* biology. Because cat intestinal organoids are difficult to culture (21, 22), the goal of this study was to understand the cell type-specific effects of LA on parasites grown in human intestinal cells versus human fibroblasts.

Aside from intestinal organoids, our knowledge of LA’s effects on *T. gondii* are largely limited to human foreskin fibroblasts (HFFs). In HFFs, LA inhibits parasite growth at physiological concentrations of 100-300 µM (23, 24, 25). However, HFFs have lower metabolic activity than gut epithelial cells, making them a poor choice for modeling fatty acid metabolism and parasite sexual development. Instead, we tested the human colon cancer cell line Caco-2. Though Caco-2 cells have altered metabolism due to their cancerous state, we selected them for this study because they are commonly used to model small intestinal absorption of drugs and nutrients and they are experimentally tractable (26, 27). We found that LA is much less potent in Caco-2 cells than in HFFs. Our data suggest that highly metabolic and abundant Caco-2s sequester LA away from *T. gondii* tachyzoites, thereby preventing LA-induced lipotoxicity. Our work highlights important differences in cell type-specific lipid metabolism that will help improve cell culture models for *T. gondii* sexual development.

## Materials and Methods

### Parasite and host cell culture

A549, Caco-2, and human foreskin fibroblast (HFF) cells were grown in a humidified 37°C incubator with 5% CO_2_. All cells were cultured in Dulbecco’s modified Eagle’s medium (DMEM, Gibco) supplemented with 10% fetal bovine serum, 2 mM L-glutamine or GlutaMAX, 10 mM HEPES, and 1% penicillin-streptomycin. Cultures considered “old” for the purposes of Figure 4 were grown for an additional 2 weeks post-confluence, with media changes every 3-5 days for A549 and Caco-2 cells and once a week for HFFs. *T. gondii* tachyzoites were grown in HFFs under the same conditions. Parasites included in this work include: Type I, RH and RH mCherry; Type II, Pru, Pru Cre mCherry (a kind gift from Anita Koshy’s laboratory), ME49 mCherry (28), and ME49 Δhpt luciferase (a kind gift from Jeroen Saeij’s laboratory); Type III, VEG; Type I/III, EGS DoubleCat (a kind gift from Louis Weiss’s laboratory, 29). To avoid biological artifacts from prolonged passage in tissue culture, Type II and III tachyzoites were maintained up to 20 passages in HFFs and then passaged once *in vivo* through mice to obtain “passage 0” bradyzoites from brain cysts. “Passage 0” ME49 mCherry and ME49 Δhpt luciferase tachyzoites were derived from the brain cysts of oocyst-infected mice.

### Animal infections for parasite propagation

Mice were treated in compliance with the guidelines set by the Institutional Animal Care and Use Committee (IACUC) of the University of Wisconsin School of Medicine and Public Health (protocol #M005217). The institution adheres to the regulations and guidelines set by the National Research Council.

### Fatty acid and drug preparations

Data in Figure 1b and part of Figure 4a were collected using an aqueous ∼3.5 mM solution of linoleic acid conjugated to 10% w/v BSA purchased from Sigma (cat no. L9530), approximately a 2:1 FA:BSA ratio. For remaining experiments, we prepared our own fatty acid mixtures by conjugating them to a stock solution of100 mg/mL (10% w/v) BSA in PBS (Sigma A9418-10G). Fatty acids were purchased from Nu-Chek Prep (linoleic acid, U-59-A; and oleic acid, U-46-A). Solutions were prepared by adding 2 µL linoleic acid or 1.98 µL oleic acid to 1 mL of 10% w/v BSA plus 4 µL per mL 1M NaOH. Solutions were incubated and rotated at 37°C for ∼3 hours or until oil droplets were no longer visible, to obtain a 7 mM solution of LA or OA with a FA:BSA ratio of 4.67:1. Solutions were filtered through a 0.8 µm filter, aliquoted, and stored at −20°C until the time of experiments. Dihydrochlamydocin (Cayman Cay21614-1) was dissolved in DMSO to 1 mM and stored at −80°C until the time of experiments. By necessity, 350 µM Sigma LA-BSA comprised 10% media volume. For all other experiments, BSA, OA-BSA, LA-BSA, and DHC (diluted in BSA) comprised 5% of total media volume, such that BSA was at 0.5% w/v (75 µM) across all treatments.

**Figure 1.**
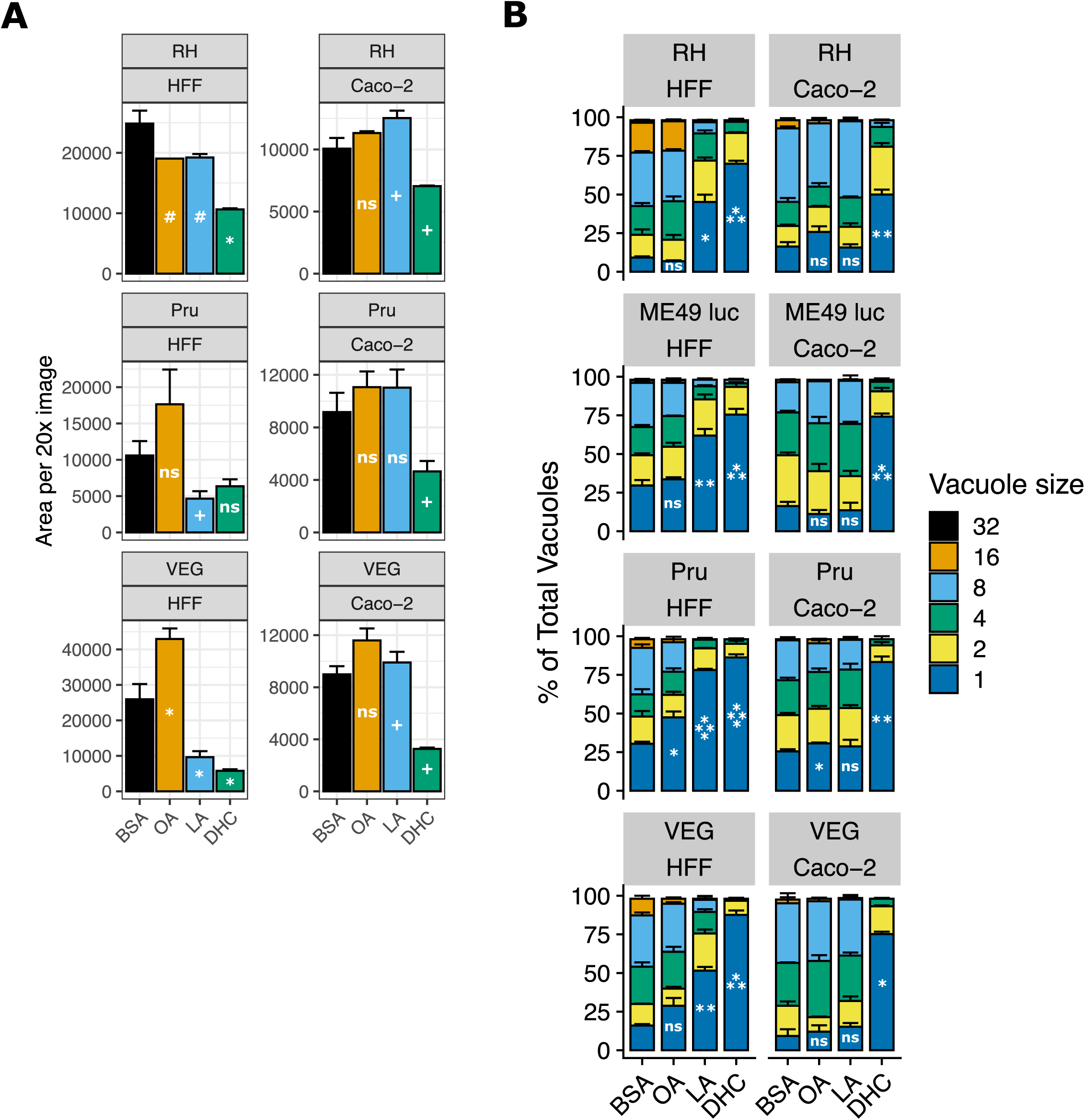
Linoleic acid slows parasite growth in HFF cells but not in Caco-2 cells. (a) *T. gondii* abundance after 4 days of treatment with 350 µM OA, 350 µM LA, 500 nM DHC, or an equal volume of BSA (final concentration 75 µM) in confluent HFFs or Caco-2 cells. After fixing and staining parasites red with immunofluorescence, 16 technical replicate images for each of 3 biological replicate wells were collected at 20x magnification on the Incucyte imaging system. Red fluorescence area was calculated for each technical replicate and averaged to obtain a value for each biological replicate. The mean of biological replicates is displayed on the y-axis +/- SE. #, p < 0.15; +, p < 0.1; *, p < 0.05 by Student’s t-test. One representative experiment of two is shown. (b) *T. gondii* replication assessed by parasitophorous vacuole (PV) size after 24 hours of treatments noted in 1a. Number of parasites per vacuole are shown as mean percentages of total PVs +/- SE. At least 100 vacuoles were counted per biological replicate. One representative experiment of two is shown, with 3 biological replicate wells per condition. *, p < 0.05; **, p < 0.01; ***, p < 0.001; ****, p < 0.0001 by Student’s t-test for percent of single-parasite vacuoles.

### Replication assays

Replication assays were performed by seeding HFF and Caco-2 cells on sterile glass coverslips in 24 well plates. Cell monolayers were infected with 50,000 parasites of VEG, ME49 Δhpt luciferase, Pru, and RH in quadruplicate for each condition. Cells infected with RH were allowed to invade for 30 minutes, whereas cells with VEG, ME49 Δhpt luciferase, and Pru were allowed to invade for 3-4 hours. The media was then replaced with drug media containing 75 µM BSA, 350 µM OA-BSA, 350 µM LA-BSA, 500 nM DHC and 75 µM BSA. At 24 hours post-treatment, the plates were fixed with 4% paraformaldehyde, permeabilized for 5 minutes with 0.2% Triton-X-100, and blocked for 1 hour at room temperature or overnight at 4°C in 3% BSA. Primary antibodies were incubated overnight at 4°C. Primaries used: 1:750 rabbit polyclonal anti-*T. gondii* (Thermo Scientific PA17252) with 1:750 goat anti-rabbit Alexa 594 secondary antibody (Thermo Scientific A-11005), or 1:750 mouse anti-*T. gondii* polyclonal (in-house) with 1:750 goat anti-mouse Alexa 594 secondary antibody (Thermo Scientific 11-005). Secondary antibodies and DAPI were incubated 1 hour at RT. Coverslips were mounted onto slides with Vectashield (Vector Laboratories H-1700-2) and the number of parasites per vacuole were counted for each condition (n=3-4 coverslips per condition). Values were imported into Excel and then RStudio for statistical analysis and figure preparation.

### SAG1 staining

For experiments investigating membrane integrity with the anti-SAG1 antibody, we prepared parasites with the same treatments and conditions noted for replication assays up to primary antibody addition. For staining, we used a monoclonal anti-SAG1 from a 1:1000 dilution of cell supernatant from the DG52 mouse hybridoma, a kind gift from John Boothroyd’s laboratory, combined with a goat anti-mouse-Alexa 488 secondary antibody at 1:750 (Thermo Scientific A-11001). Coverslips were mounted onto slides with Vectashield and imaged on a confocal microscope (ZeissLSM 800 Laser Scanning Microscope).

### High throughput growth and differentiation assays

We used an Incucyte Live Cell Analysis system (Sartorius) to assess tachyzoite growth rates under different conditions. The Incucyte was housed in a humidified 37°C incubator with 5% CO_2_. Host cells were grown to confluence in 48- or 96-well plates and infected with 2500 or 1000 tachyzoites per well, respectively. Fluorescent parasite strains were then housed in the Incucyte for the duration of the experiment and imaged every 12-48 hours. Unless specified otherwise, 0-hour reads correspond to the time immediately following drug addition or a different culture medium. Non-fluorescent parasites were grown 3-6 days in their home incubator, fixed, stained as noted in Replication Assays, and imaged once. Culture media was changed every 2-3 days for fatty acid and drug treatments, and weekly for bradyzoite switch media. Because the HDAC inhibitor DHC is such a potent inhibitor of parasite growth, it was only given to parasites for 24 hours, after which media was replaced with media containing only BSA. Parasites growing at 33°C were kept in those conditions except during imaging (approximately 1 hour every 24-48 hours). Images were obtained with at 20x magnification using phase, green, and red fluorescent channels. Automated image analysis settings were manually optimized for each plate to exclude background host fluorescence (which varied by host cell type) without excluding parasite fluorescence (which varied by strain and experiment). Out-of-focus images were excluded, as were images with large chunks of autofluorescent debris or non-confluent areas of the host monolayer. Resulting per-image data for parasite counts and total fluorescent area were exported into Microsoft Excel and then RStudio for statistical analysis and figure preparation.

### Lipid droplet quantitation

Host cells were grown to confluence in 24-well plates on sterile glass coverslips. Cells were infected with 100,000 ME49 Δhpt luciferase tachyzoites and allowed to invade and replicate for 24 hours. Cultures were then treated with 75µM BSA, 350 µM OA-BSA, or 350 µM LA-BSA for 24 hours. Cells with fixed with 4% PFA for 20 minutes at room temperature (RT), washed with PBS, permeabilized for 5 minutes at RT with 0.2% Triton-X-100, blocked for 1 hour at RT with 5% BSA, and stained red using the antibodies and conditions noted in Replication Assays. After the secondary antibody/DAPI stain, we stained samples with a 1:2500 dilution (in PBS) of a DMSO stock solution of 5 mM BODIPY 493/503 (Thermo Scientific D3922) for 20-30 minutes at RT. Coverslips were mounted with Vectashield and images collected on a confocal microscope (ZeissLSM 800 Laser Scanning Microscope) with the 63x oil objective, using Z-stacks to ensure lipid droplets were inside parasites. Representative images were collected with the 100x oil objective.

### Linoleic acid quantitation

HFFs and Caco-2s were grown to confluence in T25 flasks and treated with 75µM or 350 µM LA-BSA in quadruplicate for 24 hours. One milliliter of supernatant was collected from each flask and quickly mixed on ice with N-butanol extraction solvent at a 6/2/1 v/v/v ratio of N-butanol/acetonitrile/cell supernatant. Samples were vortexed for 10 seconds, incubated on ice 10 minutes, and spun at 4°C for 2 minutes at 14,000xg. 80-800µL of supernatant was transferred to PTFE-lined glass screwtop vials (Sigma 27000), avoiding pelleted debris. Samples were evaporated under nitrogen gas until dry and stored at −20°C until time of analysis. Ultra high pressure liquid chromatography–high resolution mass spectrometry (UHPLC– HRMS) data were acquired using a Thermo Scientific Q Exactive Orbitrap mass spectrometer coupled to a Vanquish UHPLC operated in negative ionization mode. All solvents used were of spectroscopic grade. Each sample was diluted with 100% methanol in concentration of 1 mg/mL and filtered with 0.2 µm syringe filter. A Waters XBridge BEH-C18 column (2.1 × 100 mm, 1.7 μm) was used with acetonitrile (0.1% formic acid) and water (0.1% formic acid) as solvents at a flow rate of 0.2 mL/min. The screening gradient method for the samples is as follow: Starting at 55% organic hold for 1 min, followed by a linear increase to 98% organic over 18 min, holding at 98% organic for 2 min, for a total of 21 min. A quantity of 10 μl of each sample was injected into the system for the analysis. Purchased linoleic acid (Sigma-Aldrich L1376-10G) was used as standard. For the quantification, standard curve for linoleic acid was calculated based on intensities from 5 different concentrations (25, 12.5, 6.25, 3.125, and 1.5625 ppm).

### Lipidomics

Following supernatant removal for LA quantitation (above section), cell pellets were prepared for lipidomics as follows. Supernatant was removed from flasks, and cell monolayers washed twice with warm PBS. Monolayers were scraped using plastic cell scrapers, spun at 2500xg for 15 seconds, washed once more with PBS, and spun again. Resulting pellets were frozen at −80°C until time of analysis. Each pooled sample or individual sample was analyzed separately in both positive and negative ionization modes. This was to permit different dilution factors to be employed for each mode to maximize lipid annotations. In this sample set it was experimentally determined a 40x dilution was appropriate for the positive mode and a 2x dilution for the negative mode. Downstream processing of data was conducted independently for each ionization mode. Lipid libraries were obtained by iterative MS/MS analysis of a pooled sample representing the treatment conditions and processed using Lipid Annotator (Agilent) software. This produced 4620 and 4517 features, and 375 and 560 identified lipids in positive and negative modes respectively. Individual sample lipid quantifications were obtained by MS1 analysis and processing with Profinder (Agilent) software utilizing the lipid library for that ionization mode.

#### Sample extraction

All solutions were pre-chilled on ice. Lipids were extracted in a solution of 250 µL PBS, 225 µL methanol containing internal standards (Avanti SPLASH Lipid Mix, Cayman: Carnitine #35459, Ceramide #22788, Linoleic Acid #9002193 and EOS #24423 at 10 µL per sample) and 750 µL MTBE (methyl tert-butyl ether). The sample was homogenized in three 30 s cycles alternated with a 5 minute rest on ice using a Qiagen TissueLyzer. The final rest on ice being 15 minutes. After centrifugation at 16,000 x g for 5 minutes at 4 °C, 500 µL of the upper phases were collected and evaporated until dry under a Savant speedvac concentrator. Lipid samples were reconstituted in 150 µL isopropyl alcohol and transferred to an LC-MS vial with insert (Agilent 5182-0554 and 5183-2086) for analysis. Concurrently run were a process blank sample and pooled quality control (QC) sample, prepared by taking equal volumes (∼10 µL) from each sample after final resuspension.

#### LC-MS methods

Lipid extracts were separated on an Agilent InfinityLab Poroshell 120 EC-C18 1.9 µm 2.1 x 50 mm column maintained at 50°C connected to an Agilent HiP 1290 Multisampler, Agilent 1290 Infinity II binary pump, and column compartment connected to an Agilent 6546 Accurate Mass Q-TOF dual ESI mass spectrometer. For positive mode, the source gas temperature was set to 250 °C, with a gas flow of 12 L/min and a nebulizer pressure of 35 psig. VCap voltage was set at 4000 V, fragmentor at 145 V, skimmer at 45 V and Octopole RF peak at 750 V. For negative mode, the source gas temperature was set to 350 °C, with a drying gas flow of 12 L/min and a nebulizer pressure of 25 psig. VCap voltage was set at 5000 V, fragmentor at 200 V, skimmer at 45 V and Octopole RF peak at 750 V. Reference masses in positive mode (*m/z* 121.0509 and 922.0098) and negative mode (*m/z* 1033.988, 966.0007, and 112.9856) were delivered to the second emitter in the dual ESI source by isocratic pump at 15uL/min. Samples were analyzed in a randomized order in both positive and negative ionization modes in separate experiments acquiring with the scan range m/z 100 – 1500. Mobile phase A consisted of ACN:H_2_O (60:40 v/v) containing 10 mM ammonium formate and 0.1% formic acid, and mobile phase B consisted of IPA:ACN:H_2_O (90:9:1 v/v) containing 10 mM ammonium formate and 0.1% formic acid. The chromatography gradient for both positive and negative modes started at 15% mobile phase B then increased to 30% B over 2.4 min, it then increased to 48% B from 2.4 – 3.0 min, then increased to 82% B from 3 – 13.2 min, then increased to 99% B from 13.2 – 13.8 min where it was held until 15.4 min and then returned to the initial conditioned and equilibrated for 4 min. Flow is 0.5 mL/min throughout, injection volumes were 1µL for positive and 5 µL negative mode. Tandem mass spectrometry was conducted using the same LC gradient at collision energy of 25 V.

#### Data analysis and pretreatment

QC samples and blanks were injected throughout the sample queue to ensure the reliability of acquired lipidomics data. Results from LC-MS experiments were collected using Agilent Mass Hunter (MH) Workstation Data Acquisition. Briefly, a pooled lipid extract comprised of an aliquot from each sample was analyzed in MS/MS mode. From this data, a lipid library was created using Lipid Annotator (Agilent Technologies, Inc) for positive and negative ion data, incorporating lipid identity (with either enumerated acyl chain composition or a sum composition), m/z value, and retention time. Data for all individual samples was collected in MS mode. Profinder (Agilent Technologies, Inc.) was used to perform retention time alignment between samples and to extract peak areas from each sample, for each lipid present in the library generated by Lipid Annotator. The results from Profinder were exported to .csv files for positive and negative ionization modes, reported by area. Area values were imported into RStudio for statistical analysis and figure preparation.

### Mitomycin C inactivation

Mitomycin C (R&D Systems 3258/10) was dissolved in DMSO to 10 mg/mL and stored at −20°C until time of experiments. It maintained effectiveness for up to 6 months at −20°C. For mitomycin C experiments in Figure 4, Caco-2 cells were grown to 95% confluence and treated with 10 µg/mL mitomycin C-containing media for 2-3 hours. After 2 PBS washes, cells were given normal media again until 3-4 days post-MMC treatment. At that point, cells were infected.

## Results

### Linoleic acid slows parasite growth in HFFs but not Caco-2 cells

To determine how LA affects *T. gondii* in Caco-2 cells versus HFFs, we first assessed how LA affects parasite growth using an Incucyte live cell imaging system. Briefly, we grew *T. gondii* tachyzoites of the three major type strains: RH (type I), Pru (type II), and VEG (type III) in the presence of 350 µM LA or oleic acid (OA, C18:1) for 3-4 days. We then fixed, stained, and quantitatively imaged parasites using the Incucyte. LA-treated VEG parasites were significantly less abundant in HFFs than bovine serum albumin (BSA)-treated controls, with similar trends for RH and Pru (Fig. 1b, Fig. S1a). In HFFs, OA either had no effect on parasites or enhanced their growth. Both fatty acids tended to enhance growth in Caco-2 cells.

As a positive control for growth inhibition, we used dihydrochlamydocin (DHC), an HDAC inhibitor with potent anti-*T. gondii* activity (Bougdour et al. 2009). 500nM DHC inhibited parasite growth in both cell types, highlighting that only LA has a cell-specific phenotype. Soon after these initial experiments, we changed fatty acid vendors due to supply chain issues. When we repeated the Incucyte experiments, we noted that LA purchased from Nu-Chek Prep was more inhibitory than LA purchased from Sigma (Fig. S1b) with higher bioavailability, as assayed by increased lipid droplet formation in HFFs (Fig. S1c). For biological and logistical reasons, we performed subsequent experiments with Nu-Chek Prep fatty acids unless indicated otherwise.

To determine the short-term effects of LA on *T. gondii* replication, we assessed parasitophorous vacuole (PV) size after 24 hours of fatty acid treatment. As expected given previous studies in fibroblasts (24, 25), LA-treated parasites had significantly fewer parasites per vacuole in HFFs relative to the BSA treatment (Fig. 1b, Fig. S1d). LA did not slow parasite replication in Caco-2 cells except in one experimental replicate for ME49 Δhpt luciferase parasites (Fig. S1d). OA only slightly slowed parasite replication, if at all. DHC inhibited parasite replication to a similar degree in all parasite strains and in both host cells, again highlighting the cell specific-nature of LA inhibition.

### Linoleic acid is parasiticidal

Like other antimicrobials, antiparasitics can act as either parasiticidal compounds that kill parasites or as parasitistatic compounds that do not kill parasites but halt their growth. To determine which mode of action fit LA, we monitored live parasite populations over time using mCherry-expressing strains of *T. gondii* and the Incucyte system (Fig. 2). At 350 µM LA, populations of both RH and ME49 continued to grow for 24-48 hours post-LA treatment before decreasing to below their initial abundance (Fig. 2a, Fig. S2a,b). For RH-mCherry, we were also able to track parasite expansion and contraction at the level of individual vacuoles (Fig. 2b). In keeping with the population-level pattern, we found that most vacuoles grew for 24-48 hours before slowly decreasing in size and disappearing. Lower doses of LA did not kill RH-mCherry parasites, but they grew more slowly than BSA-treated controls.

**Figure 2.**
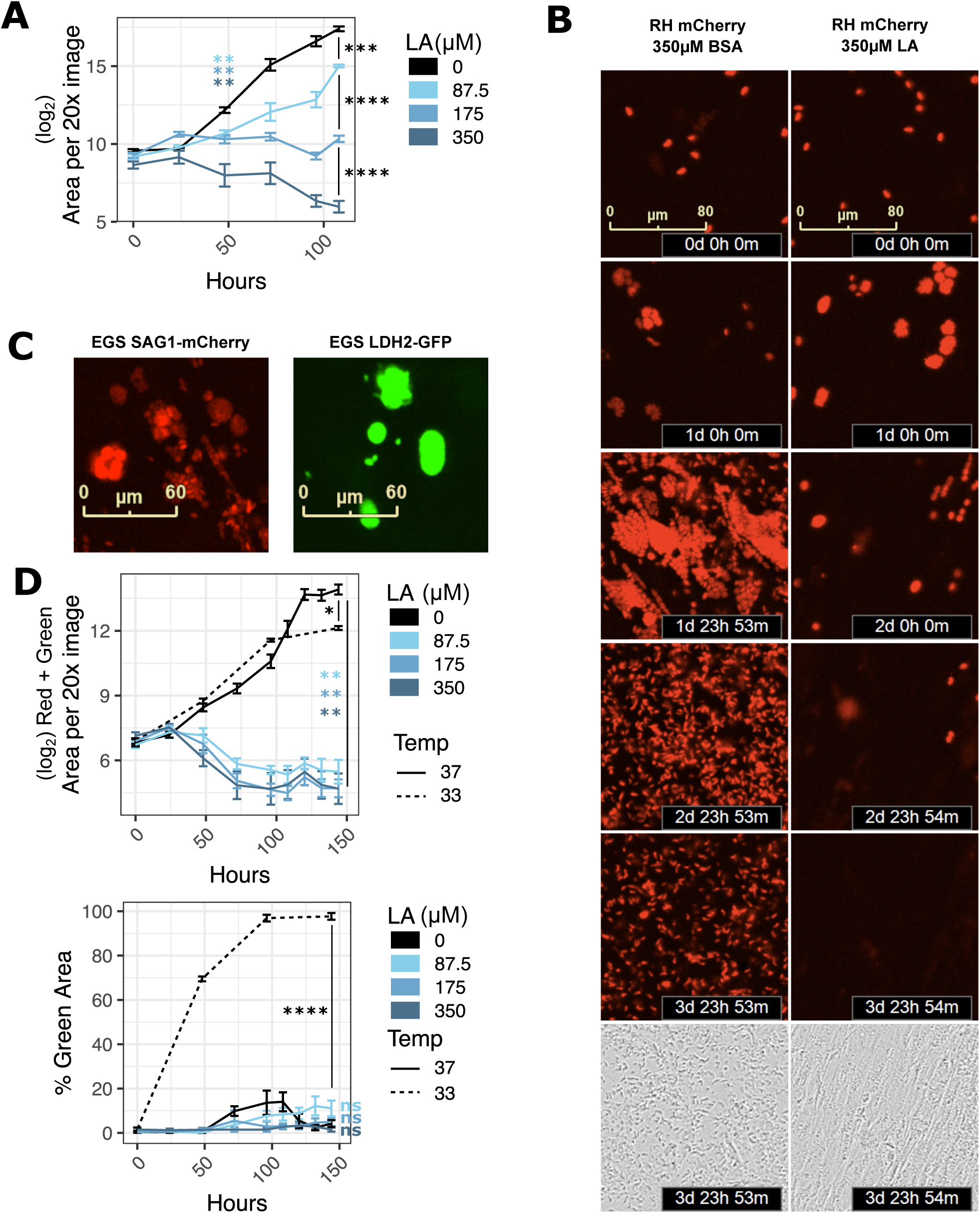
Linoleic acid is parasiticidal in HFFs. (a) RH-mCherry *T. gondii* abundance after 4.5 days of treatment with LA at indicated doses in confluent HFFs. Time 0 images were collected immediately following addition of LA at indicated micromolar doses. 16 technical replicate images for each of 4 biological replicate wells were collected at 20x magnification on the Incucyte imaging system every 12-24 hours. Red fluorescence area was calculated for each technical replicate and averaged to obtain values for each biological replicate. The mean of the log_2_-transformed biological replicate values is displayed on the y-axis +/- SE. **, p < 0.01; ***, p < 0.001; ****, p < 0.0001 by Student’s t-test. (b) Red fluorescent images from 2a are shown at 24-hour increments up to 4 days post-treatment, and phase images from 2a are shown at 4 days post-treatment. (c) Representative images of tachyzoite-stage (left, red) and bradyzoite-stage (right, green) EGS DoubleCat *T. gondii*, Incucyte imaging system, 20x magnification. (d) EGS DoubleCat *T. gondii* abundance after 6 days of treatment with LA at indicated doses (µM) in confluent HFFs. 37°C = tachyzoite-promoting conditions with CO_2_ and neutral pH. 33°C = bradyzoite-promoting conditions without CO_2_ and pH 8.0 growth media. 16 technical replicate images for each of 4 biological replicate wells were collected at 20x magnification on the Incucyte imaging system every 12-24 hours. Areas of either the sum of red and green (top) or green divided by the sum of red and green (bottom) were computed and averaged by biological replicate. Either log_2_-transformed means +/- SE (top) or mean proportions +/- SE (bottom) of biological replicates are displayed on the y-axis. *, p < 0.05; **, p < 0.01; ****, p < 0.0001 by Student’s t-test.

A very small percentage of parasite vacuoles persisted in spite of LA treatment, suggesting that some parasites are not killed by LA. Type II and III strains of *T. gondii* survive some environmental stresses by differentiating into slow-growing bradyzoites. To see if parasites survive LA-induced stress by converting to bradyzoites, we tested a highly cystogenic Type I/III EGS strain that was engineered as a dual tachyzoite-bradyzoite reporter strain (29). This strain, called EGS DoubleCat, fluoresces red (mCherry) under the control of the tachyzoite-specific SAG1 promoter and green (GFP) under the control of the bradyzoite-specific LDH2 promoter. To confirm that EGS DoubleCat fluorescence was compatible with the Incucyte system, we infected confluent HFFs with DoubleCat tachyzoites for 2 hours and switched to high pH bradyzoite induction media. We then grew parasites at 33°C in a humidified incubator without carbon dioxide. Within 4 days, >=90% of parasites switched from red to green fluorescence, indicating bradyzoite differentiation (Fig. 2c right, Fig. 2d). Consistent with bradyzoite behavior, total vacuole counts rose for one week post-treatment while maintaining >=90% GFP positivity and leaving the host monolayer intact (Fig. S2c). In contrast, nearly all parasites grown in regular media at 37°C maintained red fluorescence (Fig. 2c left, Fig. 2d), with a small fraction either switching to green or expressing neither red nor green.

Having confirmed the EGS DoubleCat strain’s ability to differentiate and be detected using the Incucyte system, we next tested whether LA induces bradyzoite formation. In contrast to high pH media and 33°C conditions, LA at 350, 175, or 87.5 µM did not increase the percentage of GFP-expressing parasites relative to the BSA treatment (Fig. 2d). Instead, overall parasite abundance decreased with a very small fraction of vacuoles persisting as in the RH and ME49 strains. These data suggested that LA lacks the capacity to induce bradyzoite formation in HFFs.

### Parasites accumulate lipid droplets and membrane aberrations following linoleic acid treatment

While performing replication assays, we noticed that LA-treated ME49 Δhpt luciferase parasites in HFFs tended to be smaller and slightly misshapen (Fig. 3a). Similar findings were reported by Nolan *et al.* (24) in a model of OA-induced lipotoxicity. Previous studies of *T. gondii* lipotoxicity also reported unusual membrane protrusions and extracellular cytoplasmic contents (24, 30). We noted similar SAG1-positive membrane protrusions and extracellular mCherry in RH-mCherry parasites that were exclusive to LA treatment in HFFs (Fig. 3b). In both parasite strains and both host cell types, DHC treatment resulted in enlarged, rounded vacuoles. Collectively, these observations pointed to lipotoxicity as a possible mechanism for parasite death in LA-treated HFFs and a different mechanism for DHC-induced parasite death.

**Figure 3.**
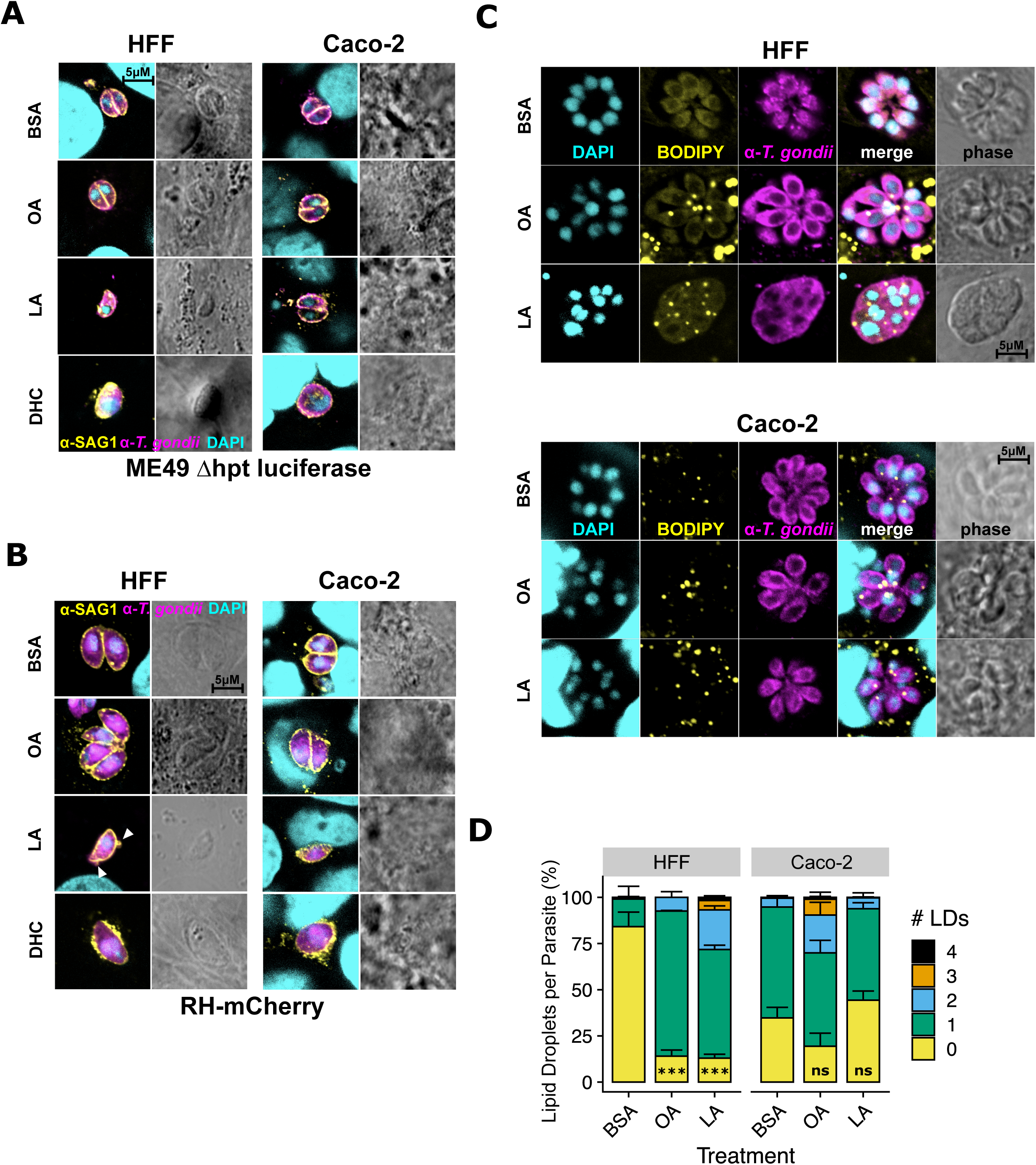
In HFFs, linoleic acid disrupts parasite membranes and increases lipid droplets. (a,b) Representative immunofluorescence microscopy images of ME49 Δhpt luciferase (a) and RH-mCherry (b) parasite vacuoles after 24 hours of treatment with 350 µM OA, 350 µM LA, 500 nM DHC, or an equal volume of BSA (final concentration 75 µM) in confluent HFFs or Caco-2 cells. Monolayers were fixed with formaldehyde and stained with a polyclonal anti-*T. gondii* antibody (ME49 Δhpt luciferase strain, magenta) and a mouse monoclonal anti-SAG1 antibody (ME49 and RH-mCherry, yellow), then counterstained with DAPI (cyan). White arrowheads point to membrane disruptions. (c) Lipid droplet formation in *T. gondii* vacuoles. ME49 Δhpt luciferase tachyzoites were treated as in 3a except parasites were allowed to invade and replicate for 24 hours prior to treatment. Magenta = *T. gondii* polyclonal antibody, yellow = lipid droplets, BODIPY 493/503, cyan = DAPI. Scalebars are applicable to all photos, as images all images were collected at the same magnification. (d) Lipid droplets per parasite from 3c were counted and plotted as percent of total parasites +/- SE. At least 100 parasites were counted for each of 3 biological replicate wells. One representative experiment of two is shown. ***, p < 0.001 by Student’s t-test.

**Figure 4.**
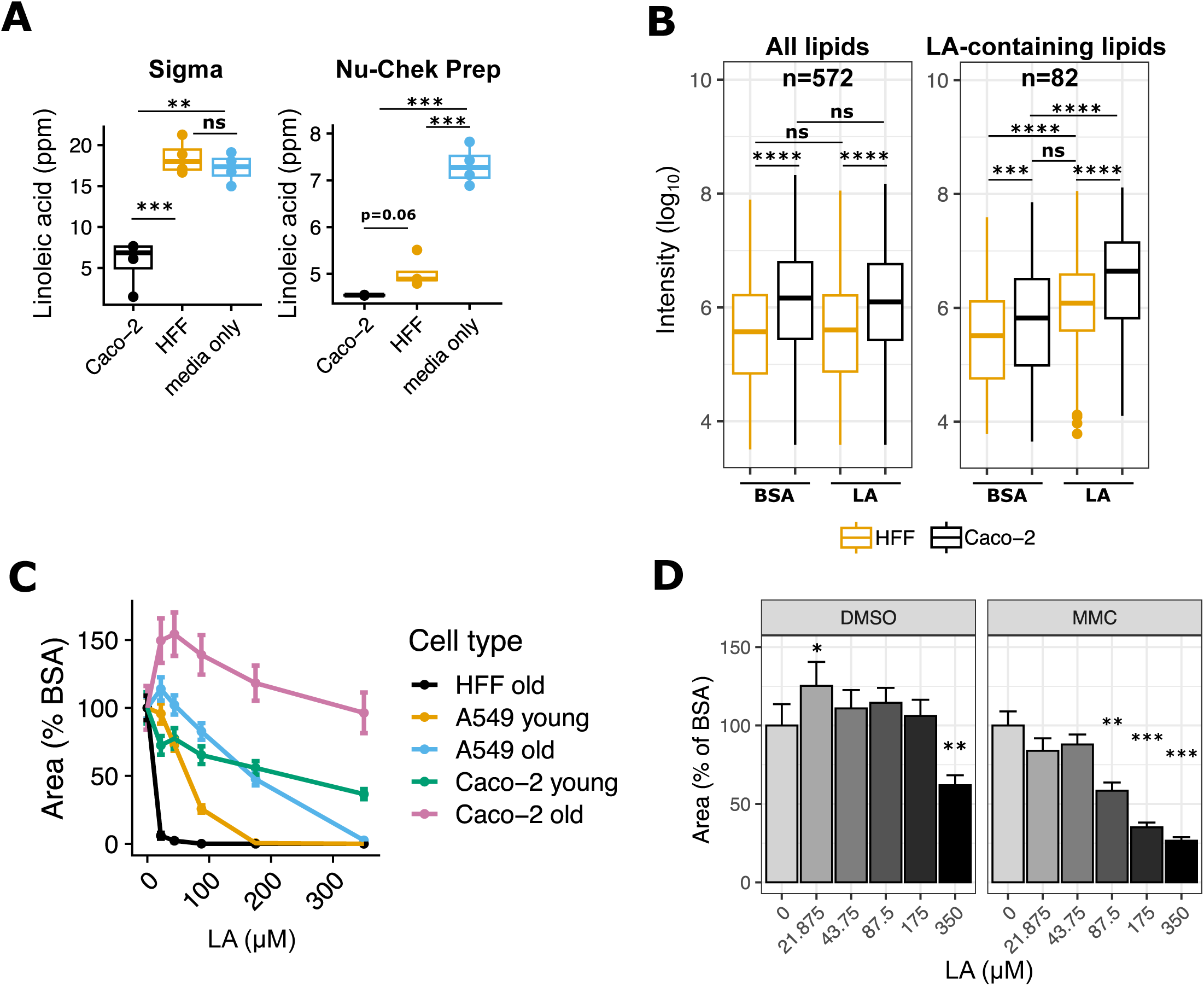
High linoleic acid metabolism in Caco-2 cells controls parasite load. (a) LA quantitation in HFF and Caco-2 supernatants, 24 hours post-treatment with 350 µM LA. Values (n=4 per condition, shown as dots) are expressed in parts per million (ppm). Two preparations of LA were tested in separate experiments on the same mass spectrometry instrument: Sigma (left) and Nu-Chek Prep (right). (b) Nu-Chek Prep LA-treated host cell pellets were sent for lipidomics analysis (n=4 per condition). Raw metabolite intensities were log_10_-transformed. Individual dots represent outliers. ***, p < 0.001; ****, p < 0.0001 by Student’s t-test. (c) Old and young cultures of Caco-2 and A549 cells and HFFs (see Methods) were infected with Pru Cre mCherry tachyzoites and grown in the presence of indicated doses of LA for 8 days. 16 technical replicate images for 4 biological replicate wells per condition were collected at 8 days post-treatment at 20x magnification on the Incucyte imaging system. Red fluorescence area was calculated for each technical replicate and averaged to obtain a value for each biological replicate. The y-axis shows the mean +/- SE of biological replicates, expressed as percent of the mean of BSA-treated control wells from the same host cell plate. (d) Pru Cre mCherry tachyzoites were grown in the presence of indicated doses of LA in Caco-2 cells that were treated with either DMSO or mitomycin C (MMC) one week prior to infection. 16 technical replicate images for 4 biological replicate wells per condition were collected at 9 days post-treatment at 20x magnification on the Incucyte imaging system. Red fluorescence area was calculated for each technical replicate and averaged to obtain a value for each biological replicate. The y-axis shows the mean +/- SE of biological replicates, expressed as percent of the mean of BSA-treated control wells. One representative experiment of two. *, p < 0.05; **, p < 0.01; ***, p < 0.001 by Student’s t-test.

Excess free fatty acids (FFAs) like OA or LA cause lipotoxicity by damaging eukaryotic membranes, organelles, or proteins (31). Organisms try to avoid FFA-induced lipotoxicity by converting FFAs into di- and triglycerides and sequestering them in lipid droplets (32). We noted a significant increase in host lipid droplet formation in both HFFs (Fig. S1c, Fig. S3a) and Caco-2s (Fig. S3a,b) in response to OA and LA treatment, suggesting both host cells respond to FFA overload by increasing lipid droplet formation. However, host monolayers remained intact even after a full week of fatty acid treatment (Fig. S3c), excluding host lipotoxicity as a cause of parasite death.

Like their mammalian hosts, *T. gondii* tachyzoites accumulate lipid droplets to avoid fatty acid overload and lipotoxicity (24). We reasoned that if LA overload causes lipotoxicity in HFF-grown parasites, those parasites would have the most lipid droplets. Indeed, parasites in LA-treated HFFs had more lipid droplets than parasites in any other treatment (Fig. 3c, Fig. 3d), except for OA-treated parasites in Caco-2s. However, given that parasites grown in Caco-2s had higher numbers of lipid droplets at baseline, the relative increase in lipid droplet formation was still highest in LA-treated HFFs. Like LA-treated parasites in HFFs in our replication assays, parasites also had disrupted membranes, as evidenced by a lack of membranes separating parasites within the same vacuole (Fig. 3c, Fig. S3a).

### Highly metabolic and abundant Caco-2 cells buffer parasites from LA toxicity

We next asked why LA-induced parasite lipotoxicity was unique to HFFs. Since parasites in LA-treated Caco-2s had fewer lipid droplets than in HFFs, we hypothesized that Caco-2s might sequester LA away from parasites by incorporating it into host lipids. To test this possibility, we measured LA uptake in each cell type by quantifying residual LA in cell supernatants after 24 hours of treatment with 350 µM LA. As expected, we found that Caco-2s took up more LA than HFFs (Fig. 4a), with some inter-experimental differences due to LA vendor. To understand which intracellular lipids incorporate LA, we sent cell pellets from the Nu-Chek Prep LA-treated samples in Fig. 4a for lipidomics analysis (Fig. 4b). We found that Caco-2 cells had higher overall levels of lipids at baseline than HFFs, including ceramides, di- and triacylglycerols, free fatty acids, sphingomyelins, and most phospholipids (Fig. S4a). LA-containing lipids were also more abundant in Caco-2s than HFFs at baseline (Fig. S4a). Both cell types responded to LA by incorporating it into all classes of lipids, but Caco-2s had the highest abundance of LA-containing lipids. These data suggest that Caco-2 cells have a higher capacity than HFFs to incorporate LA into diverse host lipids.

We next reasoned that if Caco-2s act as a sink to keep toxic LA away from parasites, then reducing the number of Caco-2s would allow more toxic LA to reach parasites. We controlled Caco-2 abundance in two ways. First, we relied on the fact that cancerous Caco-2s are not contact-inhibited and will continue to replicate the longer they are in culture. When we grew Caco-2 cells for two weeks past confluence and compared them to newly confluent cultures, the older overconfluent Caco-2 cells had higher cell counts than younger cultures (Fig. S4b). Parasite growth was inhibited by LA only in the younger Caco-2s (Fig. 4c, Fig. S4c), supporting our hypothesis that when abundant enough, Caco-2s can protect parasites from LA. To exclude the possibility that Caco-2s’ protection of parasites was just a function their cancerous state, we also looked at LA dose response curves in A549 cells, a human lung epithelial cancer cell line. Like Caco-2s, older cultures of A549s shielded parasites from LA better than young cultures. However, when controlling for cell count and LA dose, it was clear that LA was still more inhibitory to parasites in A549s than in Caco-2s (Fig. 4c, Fig. S4c, Fig. S4b). This confirmed that the effects of LA on *T. gondii* were not explained solely by host cells’ cancer states.

We applied a second method of controlling Caco-2 numbers using mitomycin C (MMC), a DNA crosslinking agent used to growth-inactivate fibroblasts for stem cell supplementation (33). Mammalian cells treated with MMC continue to be metabolically active but do not divide. After confirming that Caco-2 cells survive MMC treatment and are less abundant than DMSO-treated controls (Fig. S4d), we used these cells to compare parasite growth in the presence of LA. LA was a potent inhibitor of parasite growth in MMC-treated cells but not DMSO-treated cells (Fig. 4d, Fig. S4e), again suggesting that Caco-2 cells best protect *T. gondii* from LA-induced lipotoxicity when they are abundant.

## Discussion

We have shown that LA is parasiticidal to *T. gondii* tachyzoites in a dose- and cell-dependent manner. All parasite strains were susceptible to 350 µM LA in HFFs, which is well within the 0.2-5 mM range of LA concentrations reported for human serum (23). It is more difficult to obtain LA measurements from the small intestine, but given the abundance of LA in mammalian diets (11, 17), we can expect relatively high levels in the gut, particularly in absorptive enterocytes. Using Caco-2 cells as a proxy for enterocytes in this study, we showed that LA is much less inhibitory to parasites in gut cells than in fibroblasts or A549 lung epithelial cells. In the future, using primary intestinal epithelial cells from organoids will allow us to expand our conclusions to other gut cells.

Type I RH parasites were less susceptible to intermediate doses of LA than Type II Pru and ME49 parasites, for unknown reasons. RH parasites replicate faster and may express more fatty acid utilization or detoxification genes that prevent LA from triggering lipotoxicity. RH also possesses virulence factors that influence its interactions with host organelles that may reduce LA siphoning from the host. Type II *T. gondii* tachyzoites often differentiate into bradyzoites in the presence of stressors like high pH or arginine deprivation (34). Thus, we expected LA to induce bradyzoite formation or allow bradyzoites to persist in the presence of LA. However, our data in HFFs did not show an increase in bradyzoites following LA treatment, suggesting that parasites are unable to use differentiation to adapt to LA-induced stress. Whether LA activates bradyzoite differentiation pathways in other host cell types is yet to be determined.

Our data suggest that LA causes lipotoxicity through parasite membrane disruption, as evidenced by membrane blebs and a lack of membranes separating parasites within vacuoles. LA’s two double bonds may cause membrane disruption by increasing membrane fluidity or by increasing susceptibility to double bond oxidation. One fewer double bonds in OA may explain why OA-treated parasites appear healthy despite OA-induced lipid droplet accumulation in both host cell types. Consistent with that idea, LA and other PUFAs inhibit *Plasmodium falciparum* in a mechanism attributed to oxidation, while OA does not (35). High levels of LA could also indirectly interfere with parasite membrane integrity by modulating the biosynthesis or incorporation of other essential membrane lipids (36, 30). Future studies might investigate other mechanisms of LA-induced *T. gondii* lipotoxicity as well, such as ER stress or mitochondrial dysfunction due to excess fatty acid beta-oxidation (31). LA and other PUFAs can also influence infection and inflammation when they are enzymatically converted into oxylipins by lipoxygenases, cyclooxygenases, and soluble epoxide hydrolases (sEHs) (37, 38, 39, 40). While a soluble epoxide hydrolase (sEH) inhibitor did not influence *T. gondii* growth in our hands (data not shown), the role of oxylipins in *T. gondii* growth and development may be a promising avenue for future study.

Cultures of abundant Caco-2 cells prevented LA-induced parasite growth inhibition up to 350 µM. Even in non-abundant Caco-2 cultures, 350 µM LA was only partially inhibitory to parasites. This occurred despite high host cell uptake of LA, suggesting that Caco-2s may differ from HFFs in their ability to incorporate LA into host lipids and prevent their transfer into parasites. Our lipidomics data showed that LA was readily incorporated into all classes of Caco-2 lipids following LA treatment. While the same pattern was true for HFFs, the overall abundance of LA-containing lipids was higher in Caco-2s. Caco-2s also have more lipid droplets than HFFs following fatty acid treatment (Fig. S1, Fig. S4). These data suggest Caco-2s possess an all-around increased capacity for incorporating LA into host lipids, which is not surprising given their role as absorptive cells.

Increased LA incorporation alone does not explain Caco-2s’ protection of the parasite. *T. gondii* is a professional lipid scavenger that efficiently imports host lipids including those from lipid droplets (41, 42, 30, 43). The parasite also recruits host organelles including the ER, mitochondria, and lipid droplets to its parasitophorous vacuole membrane (PVM) to obtain nutrients. *T. gondii* therefore should be able to access Caco-2 lipids as easily as they access HFF lipids. However, it may be that the two host cell types differ in lipid trafficking or organellar association with the PVM. In support of that idea, we saw reduced parasite lipid droplet accumulation in LA-treated Caco-2s. Future work might also investigate whether Caco-2 and HFF mitochondria are recruited differently to the PVM, a process that can restrict parasite uptake of host lipids (25).

In 2018, Nolan et al. (24) found that both LA and OA inhibited *T. gondii* tachyzoites at 200 µM, with LA being the more potent inhibitor. We also found LA to be more inhibitory than OA, but we did not observe a strong inhibitory effect of OA on parasites even at 350 µM. We speculate that the difference between our studies is due to different preparations of fatty acids. Even within this study, we saw dramatic differences in LA bioavailability and parasiticidal activity depending on the LA vendor. Sigma LA was an approximately 3.5 mM aqueous solution of LA conjugated to BSA at a ratio of approximately 2:1 fatty acid:BSA. Nu-Chek Prep LA was a >99% purity oil of LA derived from safflower oil that we conjugated to BSA in-house at 7 mM and a 4.7:1 fatty acid:BSA ratio (see Methods). It is well-known that fatty acid:BSA ratios dictate bioavailability of fatty acids, with higher ratios of 6:1 mimicking pathological states (44, 45). Differences in fatty acid preparation may thus be responsible for the inter- and intra-study variation we see here.

The ultimate goal of our investigations of cell type-specific LA metabolism is to understand how the cat intestinal environment signals to *T. gondii* to enter its sexual cycle. Previous work showed that a cat-like LA metabolic environment triggers parasite commitment to sexual development (20). Our study of fatty acid metabolism in *T. gondii*-infected Caco-2 cells will help us apply those methods to cat intestinal organoids, bringing us closer to completing the entire *T. gondii* life cycle without the use of animal models.

## Data Availability

All data are contained within the article or supporting information.

## Funding and Additional Information

This work was supported by a Food Research Institute’s Summer Scholars Program (M.T.K.), National Institutes of Health National Institute of Allergy and Infectious Diseases R01AI144016-01 (L.J.K.), and a Ruth L. Kirschstein Postdoctoral Individual National Research Service Award from the National Institutes of Health National Institute of Allergy and Infectious Diseases F32 AI172084 (N.D.H.). The content is solely the responsibility of the authors and does not necessarily represent the official views of the National Institutes of Health.

## Acknowledgments

We thank Lou Weiss for the EGS DoubleCat strain, Anita Koshy for the Pru Cre mCherry strain, and David Arranz-Solís and Jeroen Saeij for the ME49 Δhpt luciferase oocysts. We thank Judith Simcox, JD Sauer, Nancy Keller, and members of the Knoll laboratory for helpful discussions. Arzu Ulu and Bruce Hammock also engaged in helpful discussion and provided the sEH inhibitor. Thank you to Michael Panas and John Boothroyd for providing the DG52 mouse hybridoma for anti-SAG1 monoclonal antibody production. We appreciate the assistance and expertise of Gregory Barrett-Wilt and Timothy Shriver from the UW-Madison Mass Spectrometry Core Facility for acquiring and analyzing lipidomics data.

---

**Figure S1. Related to Figure 1: Linoleic acid slows parasite growth in HFF cells but not in Caco-2 cells.** (a) A second replicate experiment of Fig. 1a. *T. gondii* abundance after 3 days of treatment with 350 µM OA, 350 µM LA, 500 nM DHC, or an equal volume of BSA (final concentration 75 µM) in confluent HFFs or Caco-2 cells. After fixing and staining parasites red with immunofluorescence, 16 technical replicate images for each of 3 biological replicate wells were collected at 20x magnification on the Incucyte imaging system. Red fluorescence area was calculated for each technical replicate and averaged to obtain a value for each biological replicate. The mean of biological replicates is displayed on the y-axis +/- SE. #, p < 0.15; +, p < 0.1; *, p < 0.05; **, p < 0.01; ***, p < 0.001 by Student’s t-test. (b) LA vendor influences LA potency. RH-mCherry growth was measured over time using the Incucyte, in the presence of the treatments listed in (a). (c) The Incucyte was used to count BODIPY 493/503-positive lipid droplets in HFFs in the presence of treatments listed in (a). (d) A second replicate experiment of Fig. 1b. *T. gondii* replication assessed by parasitophorous vacuole (PV) size after 24 hours of treatments noted in (a). Number of parasites per vacuole are shown as mean percentages of total PVs +/- SE. At least 100 vacuoles were counted for each of 3 biological replicate wells per condition. *, p < 0.05; **, p < 0.01; ***, p < 0.001; ****, p < 0.0001 by Student’s t-test for percent of single-parasite vacuoles.

**Figure S2. Related to Figure 2: Linoleic acid is parasiticidal.** (a, b) Additional experiments supporting conclusions from Figure 2a but with ME49-mCherry parasites. (c) A representative Incucyte phase channel image of intact HFF monolayer infected with EGS parasites after 1 week of growth at pH 8.0 in a humidified incubator without CO_2_.

**Figure S3. Related to Figure 3: In HFFs, linoleic acid disrupts parasite membranes and increases lipid droplets.** (a) 100x confocal fluorescence microscopy images of host lipid droplets in ME49 Δhpt luciferase-infected HFFs and Caco-2s following 24 hours of treatment with 350 µM OA, 350 µM LA, or an equal volume of BSA (final concentration 75 µM). Magenta = *T. gondii*, yellow = lipid droplets, cyan = DNA. (b) Incucyte-based quantification of lipid droplets in uninfected Caco-2s after 24 hours of treatment with 350 µM OA, 350 µM LA, 500 nM DHC, or an equal volume of BSA (final concentration 75 µM). (c) Incucyte images at 20x magnification of intact HFF and Caco-2 monolayers after 1 week of infection and fatty acid treatment. Red = mCherry-positive parasites.

**Figure S4. Related to Figure 4: High linoleic acid metabolism in Caco-2 cells controls parasite load.** (a) Log_10_-transformed abundances of lipids divided by lipid class. Cer = ceramides, DG = diacylglycerols, Ether = ether phospholipids, FFA = free fatty acids, LP = lysophospholipids, PC = phosphatidylcholine, PE = phosphatidylethanolamine, PG = phosphatidylglycerol, PI = phosphatidylinositol, PS = phosphatidylserine, SM = sphingomyelin, TG = triacylglycerol. All lipids (n=572) are included in the top plot. The bottom plot is restricted to LA-containing lipids (n=82). (b) Cells from the same 48-well plates used in Figure 4c were trypsinized and counted (n=6 per plate). (c) A second replicate experiment of the data shown in Fig. 4c, the cell types and abundance experiment. (d) Similar to (b), cell counts from the 48-well plates used in Fig. 4d, the mitomycin C experiment. (e) A second replicate experiment of the data shown in Fig. 4d, the mitomycin C experiment.

**Supplementary Information 1. Lipidomics data table and sample metadata.** Per-sample metabolite peak areas are reported for both positive and negative ion mode mass spectrometry runs. Sample metadata is included, as is a readme.

## Author Contributions

conceptualization: NDH, LJK

investigation: NDH, CZ, MTK, SCP

formal analysis: NDH, CZ, SCP

writing-review and editing: NDH, CZ, SCP, LJK

resources: LJK

project administration: LJK

funding acquisition: NDH, MTK, LJK

