## Supplementary figures and images for "Host cell-specific metabolism of linoleic acid controls *Toxoplasma gondii* growth in cell culture"

### Supplemental Figure 1

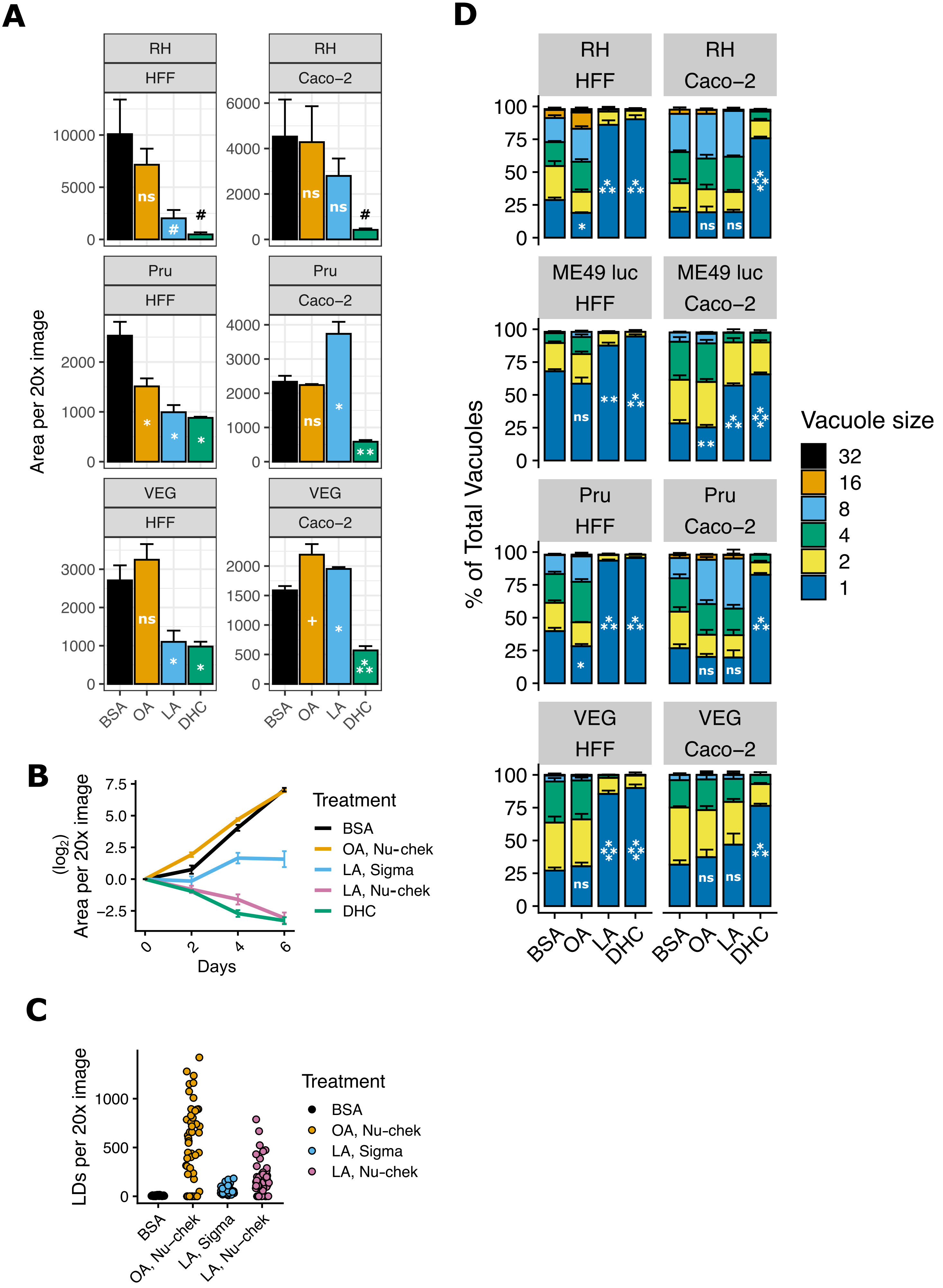

### Supplemental Figure 2

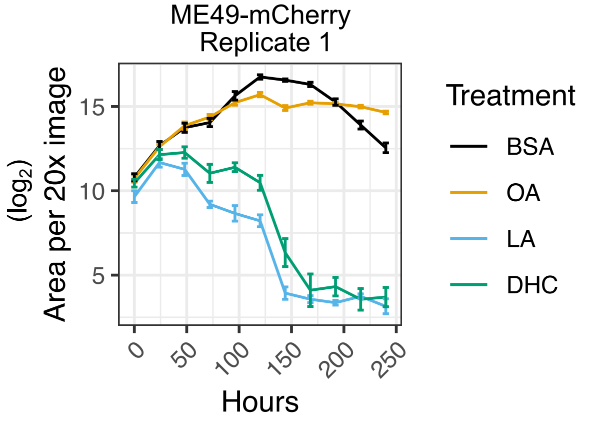

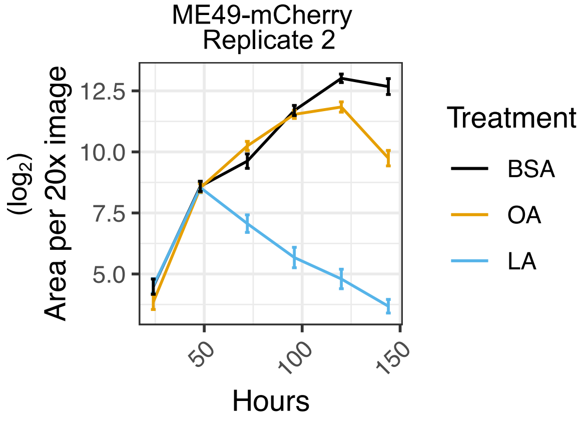

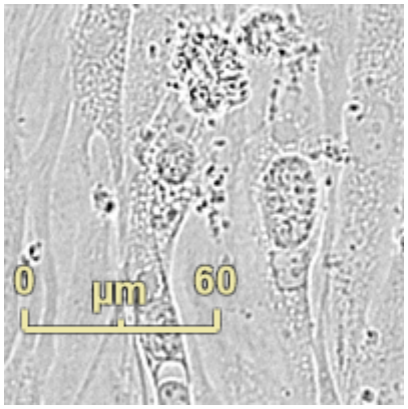

### Supplemental Figure 3

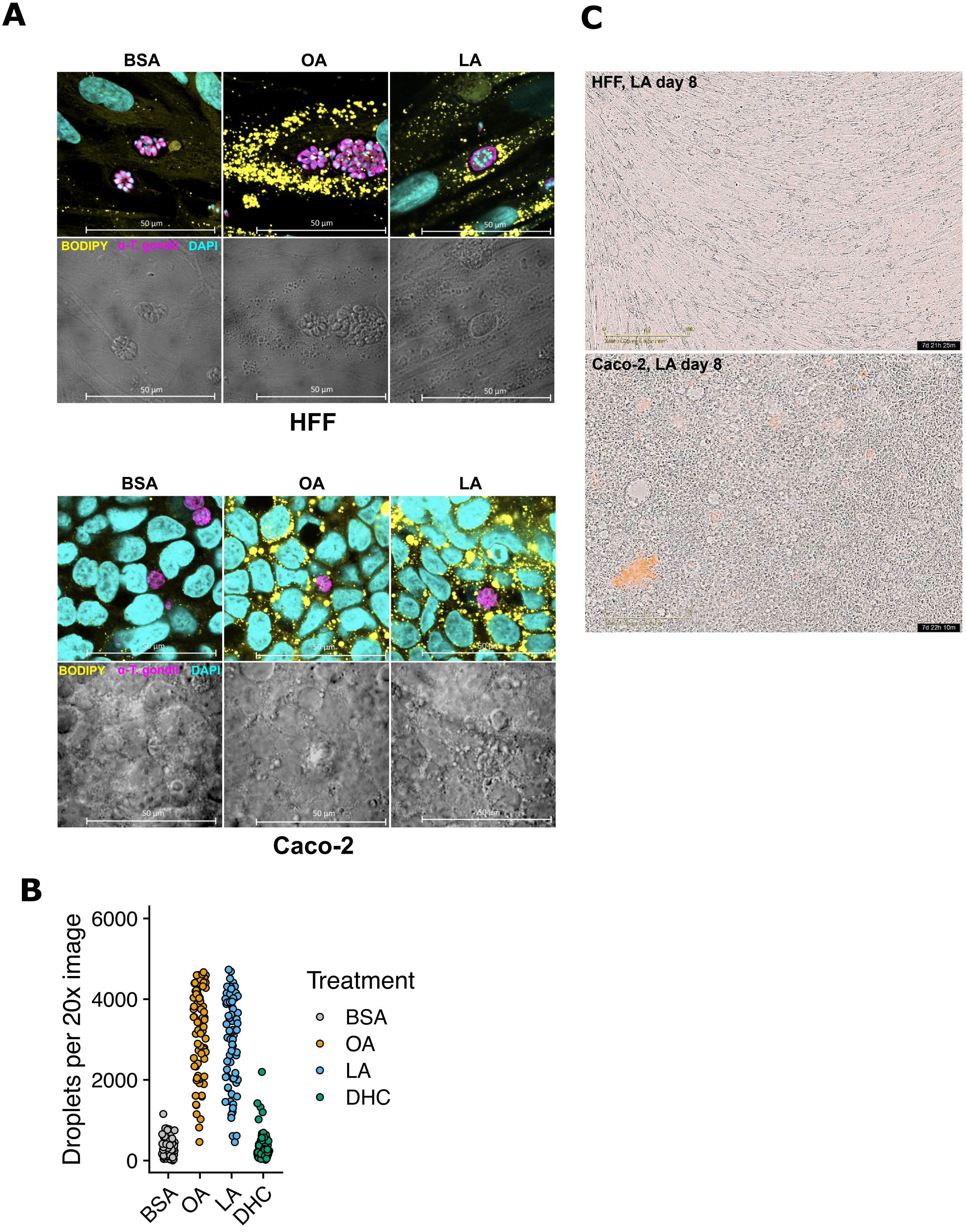

### Supplemental Figure 4

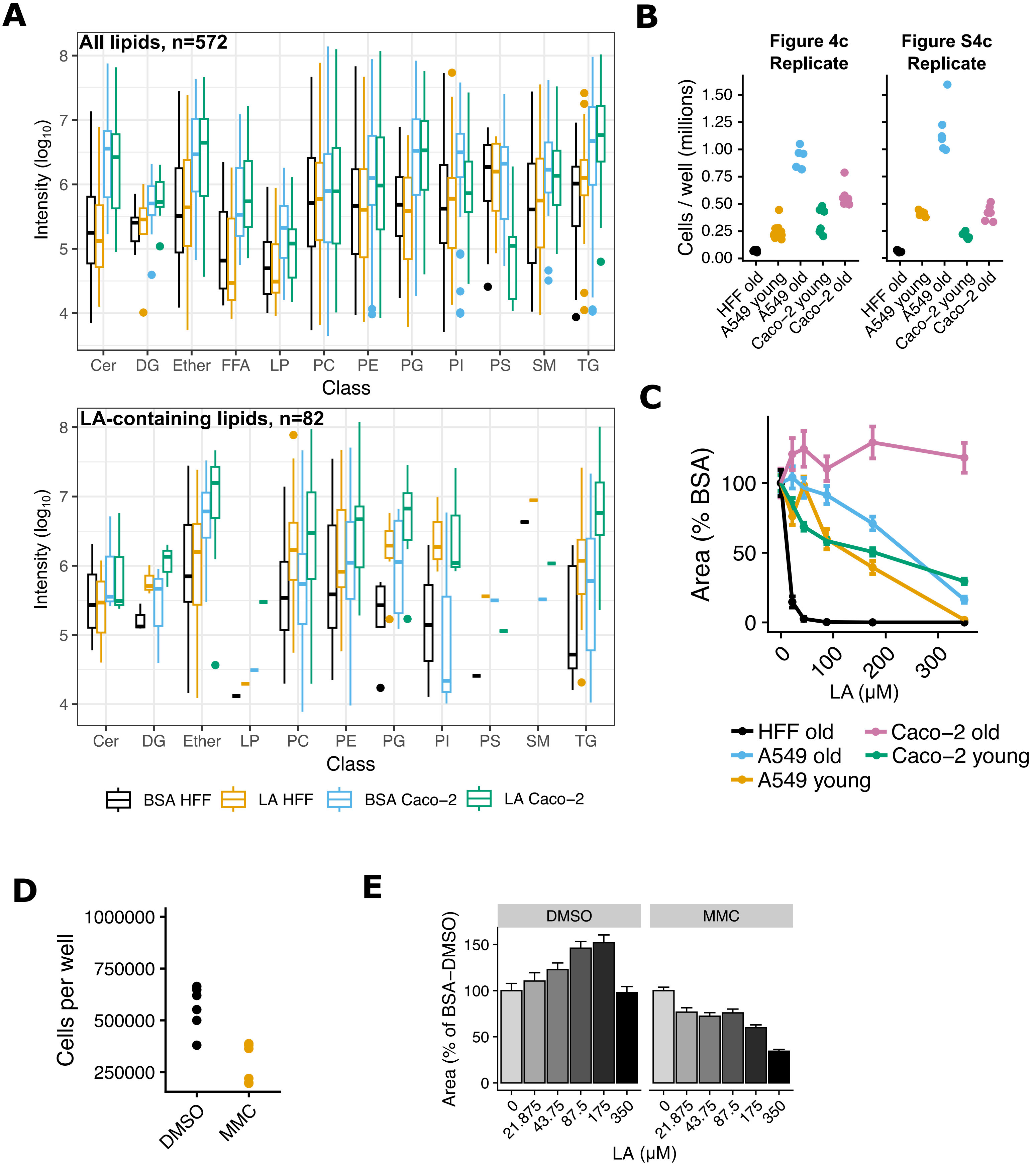
